# USH2A gene mutations in rabbits lead to progressive retinal degeneration and hearing loss

**DOI:** 10.1101/2022.01.24.476758

**Authors:** Van Phuc Nguyen, Jun Song, Diane Prieskorn, Yanxiu Li, David Dolan, Jie Xu, Jifeng Zhang, K Thiran Jayasundera, Yehoash Raphael, Y. Eugene Chen, Yannis M. Paulus, Dongshan Yang

## Abstract

Mutations in USH2A gene are responsible for the greatest proportion of hearing and vision loss among individuals with Usher Syndrome (USH) and for autosomal recessive non-syndromic retinitis pigmentosa. Mutations on USH2A exon 13 account for more than 35% of the disease causing USH2A variants including the most prevalence point mutation, c.2299delG, a frameshift mutation. The lack of a clinically relevant animal model has been a bottleneck for the development of therapeutics for USH2A related vision loss. Using CRSPR/Cas9 technology, this study establishes a rabbit line carrying an USH2A frameshift mutation on exon12 (equivalent to human USH2A Exon 13) as a novel mammalian animal model of USH2A. The bi-allelic mutant rabbits exhibit hyper reflective signals in FAF indicating RPE damage and OCT changes indicating photoreceptor degeneration as early as 4 months of age. ERG signals of both rod and cone function were reduced in the USH2A mutant rabbits starting from 7 months old and further decreased at 15-22 months old, indicating progressive retinal photoreceptor degeneration, which is further confirmed by retinal histopathology examination. ABR examination showed moderate to server hearing loss in the USH2A mutant rabbits. These results indicated that disruption of USH2A gene in rabbits is sufficient to induce hearing loss and progressive photoreceptor degeneration. To our knowledge, this is the first mammalian animal model of USH2 which closely recapitulates the phenotype of retinitis pigmentosa in human patients. This study supports the use of rabbits as a clinically relevant animal model to understand the pathogenesis and to develop novel therapeutics for Usher Syndrome.

## Introduction

Usher syndrome (USH) is an autosomal recessive genetic disorder resulting in hearing loss, progressive visual impairment and, in some types, balance issues^1^. The major ocular symptom of patients with USH is a disease called retinitis pigmentosa (RP). RP causes the light-sensing photoreceptor cells in the retina to gradually deteriorate, initially resulting in night blindness, followed by tunnel vision, and severe, permanent, progressive vision loss. More than 400,000 people are affected by USH worldwide, accounting for about 50 percent of all hereditary deaf-blindness cases^2^. USH is classified into three subtypes (I, II, and III), which are distinguished by severity and age of onset of deafness, presence or absence of vestibular dysfunction, and age at onset of RP. Among them, type II (USH2) is the most common subtype, characterized by hearing loss from birth and progressive vision loss that begins in adolescence or adulthood. USH2 may be caused by mutations in any of three genes: USH2A, GPR98, and DFNB31, with USH2A mutations being the most prevalent, present in approximately 70% of USH2 cases^3^ . Mutations in the USH2A gene are also a cause of some forms of RP without hearing loss (i.e., non-syndromic RP)^4^. More than 700 pathogenic USH2A mutations have been identified, as reported in the LOVD database (http://www.lovd.nl). Mutations in exon 13 account for approximately 35% of all USH2A cases, including the two most recurrent mutations in USH2A, c.2299delG (p. Glu767fs*21) and c.2276G>T (p. Cys759Phe) ^5^. Despite extensive research, there is no cure for USH2 yet. Hearing aids provide benefits to USH2 patients who have moderate to severe hearing loss; however, efforts to mitigate the progressive visual loss caused by RP have been disappointing. Most individuals with USH2 RP will eventually suffer from severe, progressive vision loss. Therefore, there is a pressing unmet clinical need to develop novel therapeutics for USH2.

Mouse and zebrafish models of USH2A have been developed. Unfortunately, phenotypes observed in retinas of USH2A-USH2 patients are not faithfully replicated in mouse models carrying USH2A mutations ^6–8^. USH2A knock out mice suffer from hearing loss but only manifested weak and very late onset retina degeneration phenotype. The zebrafish models exhibit early retinal degeneration phenotypes. However, their features of adult photoreceptor regeneration as well as the distance from humans may pose problems in translational studies^9^. Therefore, development of an alternative mammalian model of USH2, which more closely approximates human physiology, function, and anatomy, and importantly RP pathogenesis is of prime importance and may accelerate translating discoveries from animal models into clinical therapies and interventions for the disease.

Rabbits, compared with mice, are closer to humans in terms of phylogenesis, anatomical features, physiology, and pathophysiological responses^10–14^, and are used as a classic lab animal species to develop novel therapeutics for humans and refine medical and surgical equipment^13, 15–19^. Historically, retinal degeneration has been studied in rhodopsin Pro347Leu transgenic rabbits, a model of RP^20–31^. Recently, the emerging gene editing technology in rabbits has greatly increased their value to biomedicine, motivating our efforts to develop rabbits that carry the disease causing mutations found in human patients, as models to replicate human diseases more precisely^32–34^. In this study, we report the development of USH2A rabbits by CRISPR/Cas9. This novel model is expected to greatly facilitate both the basic and translational studies of USH.

## Materials and Methods

### Animals

New Zealand White (NZW) rabbits were purchased from Covance or Charles River. The animal maintenance, care and use procedures were reviewed and approved by the Institutional Animal Care and Use Committee (IACUC) of the University of Michigan. All procedures were carried out in accordance with the approved guidelines and were performed in accordance with the ARVO (The Association for Research in Vision and Ophthalmology) Statement for the Use of Animals in Ophthalmic and Vision Research. All efforts were made to minimize suffering. All the methods were carried out in accordance with the approved guidelines.

### Reverse transcription polymerase chain reaction (RT-PCR) and real time PCR analysis

Total RNA from retina, sclera, brain, liver, kidney, and bone marrow were isolated using the RNeasy kit (Qiagen). Reverse transcription was used to generate cDNA (SuperScript® III First-Strand Synthesis System, Thermo Fisher Scientific, 18080-05) as template for RT-PCR and real time PCR. For real time PCR analysis, samples were analyzed on a BioRad CFX Connect™ Real-Time PCR Detection System and amplification was detected using the SYBR green method (BioRad, SYBR green supermix). PCR primers are as following: RTF1: 5’-aattcaggccagtgcaagtg-3’, RTR1: 5’-gcccagaaagaggattgcag-3’; RTF2:5’-ggagaagaagagggtgtgct -3’; RTR2: 5’-gactctccactggaagctga-3’. Rabbit 18S rRNA or GAPDH expression was used as internal control. The RT-PCR products were purified and subject to Sanger sequencing.

### Scanning electron microscopy

The neuroretinas obtained from perfused rabbits were postfixed by immersion in 2.5% glutaraldehyde in 0.1 M cacodylate buffer (pH 7.3) for 2 h at room temperature. The samples were dehydrated in ethanol, dried to critical point, and fractured along a plane passing through the long axis of the photoreceptors. The fragments were mounted on aluminum stubs with double-adhesive carbon tape and examined under a scanning electron microscope.

### Histopathology

To euthanize the rabbits, euthanasia solution (Euthanasia, 0.22 mg/kg, 50 mg/mL, VetOne, ID, USA) was injected into the rabbit intravenously through the marginal ear vein. The eyeballs were harvested and fixed in Davidson’s fixative solution for 24 h. The sample was then cut into 5 mm pieces and embedded in paraffin. The sample was sectioned to a thickness of 4 µm using a Leica Autostainer XL (Leica Biosystems, Nussloch, Germany) and stained with hematoxylin and eosin (H&E). The H&E slides were observed using a Leica DM600 light microscope (Leica Biosystems, Nussloch, Germany) and the images were captured using a BF450C camera.

### CRISPR reagents

The Cas9 expression plasmid JDS246 was obtained from Addgene. Cas9 mRNA was transcribed *in vitro*, capped and polyadenylated using the T7 mScript™ Standard mRNA Production System (C-MSC100625, CELLSCRIPT, Madison, WI). Guide RNA (gRNA) was designed using CRISPOR software^35^, synthesized as chemically modified (2’-O-Methyl at 3 first and last bases, 3’ phosphorothioate bonds between first 3 and last 2 bases) single strand gRNA (sgRNA EZ Kit, Synthego). The target sequence on rbUSH2A is shown in Fig. 2A. Cas9 mRNA and sgRNA were diluted in RNase-free TE buffer (1mM Tris-Cl pH 8.0, 0.1mM EDTA), stored at -80 °C in 10 μl aliquots, and were thawed and kept on ice before microinjection.

### Rabbit genome editing

Methods of rabbit genome editing has been described previously in detail^36^. Briefly, pronuclear stage rabbit embryos were injected with approximately 2-5 pL RNase-free TE buffer (1mM Tris-Cl pH 8.0, 0.1mM EDTA) containing 150 ng/μl Cas9 mRNA, 50 ng/μl sgRNA and 50 ng/μl donor oligo. Injected embryos were washed three times in embryo culture medium, which consisted of Earle’s Balanced Salt Solution (E2888, Sigma) supplemented with non-essential amino acids (M7145, Sigma), essential amino acids (B-6766, Sigma), 1 mM L-glutamine (25030-081, Life Technologies), 0.4 mM sodium pyruvate (11360-070, Life Technologies) and 10% FBS. Twenty to thirty embryos were surgically transferred to oviducts of each synchronized recipient doe. For gRNA validation, instead of transferring to recipients, the injected embryos were washed and cultured in the medium at 38.5 °C in 5% CO2 for additional 2-3 days until they reach blastocyst stage.

### Detection of gene editing events

For gRNA *in vitro* validation, PCR products amplified the targeted USH2A gene region were purified with a PCR purification kit. The purified PCR products were mixed in a Eppendorf tube with Cas9 protein and gRNA to be tested as following: 10XNEBuffer 3.1, gRNA (30 nM final), Cas9 Nuclease, S. pyogenes (M0386S) (∼30 nM final), substrate PCR products (3 nM final), and Nuclease-free water to total reaction volume of 30 µl. The reaction solution was incubated at 37°C for 30 minutes and analyzed by gel electrophoresis.

For *in vivo* testing, injected embryos developed to blastocyst stage in culture were collected in 1.5 ul water individually and the whole genome was replicated using a REPLI-g® Mini Kit (Qiagen, Germantown, MD) following the manufacturer’s protocol. For rabbit genotyping, genomic DNA was isolated from the newborn kits ear skin. Genomic DNA was then amplified by PCR using corresponding primers: Primers for indels detection: forward (F), 5′-tctgcagtagcattgtttgtgatt-3′, reverse (R), 5′-gtcccagtctcatcacagttacaa-3′; Primers for NGS sequencing: F, 5′-agccctgccagtgtaacctc-3′, R, 5′-agtgactgagcctgctgtgttg-3′; Primers for OT1 detection: F, 5’-gaggtacaagcagggtaagaaggg-3’, R, 5’-gaatgaaacatggcctgggacct-3’; Primers for OT2 detection: F, 5’-gagagctggactggaagaggag-3’; R, 5’-agggtacttctgtgcgttcg-3’; Primers for OT3 detection: F, 5’-tcaggagtgaatcagcagatacaa-3’, R, 5’-ttcggcttattcaggaaagaaatg-3’. PCR products were purified and subjected to T7E1 assay, Sanger sequencing, and NGS (MGH CCIB DNA core). NGS data were analyzed using CRISPResso2 software^37^.

### Rabbit eye examination and imaging procedure

A comprehensive examination of the eyelids, conjunctiva, cornea, anterior chamber, iris, and lens was performed using before imaging by slit lamp bio-microscopy (SL120, Carl Zeiss, Germany). Fundus photography, fundus autofluorescence (FAF), fluorescein angiography (FA), and indocyanine green angiography (ICGA) were employed to evaluate the vascular network of the retina and choroid. Briefly, after pupil dilation, a clinical fundus camera (TRC-50EX, Topcon Corporation, Tokyo, Japan) was used to acquire fundus photography, FAF, FA, and ICGA. Fluorescein sodium (0.2mL, 10% solution) (Akorn, Lake Forest, IL, USA) and indocyanine green (0.5 mg/kg, 5 mg/mL, HUB Pharmaceuticals LLC, Patheon, Italy) were injected in the rabbit marginal ear vein. Photographs were captured immediately after injection up to 10 minutes to capture early, middle, and late phase angiography images. There was a 5-minute interval between the two kinds of angiography tests.

Spectral domain optical coherence tomography (OCT) imaging was performed as described previously^38^. Briefly, two superluminescent light emitting diodes with a center wavelength of 905 nm were used to illuminate the surface of the cornea and focused on the fundus by the rabbit eye optics. The average power of the OCT probing light was 0.8 mW. The lateral and axial resolutions are 3.8 µm and 4.0 µm, respectively. The system can achieve an imaging depth of 1.9 mm. The rabbits were put on a custom-built platform, and the eye position was adjusted under the ophthalmic lens using a CCD camera to visualize the region of interest.

### Electroretinography (ERG)

Full field ERG (ff-ERG) was performed after pupillary dilation. After 60 minutes of dark adaptation, rabbits were anesthetized. After topical anesthesia, ERG-Jet contact lens electrodes (The Electrode Store, Enumclaw, WA, USA) were applied. Corneal hydration was maintained with a 2.5% hypromellose ophthalmic demulcent solution (Akorn Inc, Lake Forest, IL, USA). A pair of reference electrodes and a ground electrode (needle electrodes, The Electrode Store) were placed subcutaneously behind the bilateral ears and in the scruff, respectively. All animal handling was done under dim red light. ERGs were recorded with a Ganzfeld configuration using the LKC UTAS 3000 electrophysiology system (LKC Technologies, Gaithersburg, MD, USA). ERG responses were amplified at 2500 gain at 0.312-500 Hz and digitized at a rate of 2000 Hz. Scotopic ERGs were recorded at a dim flash intensity of 0.01 cd.s/m2 to obtain the rod isolated ERG and at 3.0 cd.s/m2 to obtain the combined rod-cone ERG. After 10 minutes of light adaptation to a white 32 cd/m2 rod suppressing background, photopic ERGs were recorded at a flash intensity of 3.0 cd.s/m2. For ERG analyses, the a-wave amplitude was measured from the pre-stimulus baseline to the trough of the a-wave, and the implicit time of the a-wave was measured from flash onset to the trough of the a-wave. The b-wave amplitude was measured from the trough of the a-wave to the peak of the b-wave, and the b-wave implicit time was measured from flash onset to the peak of the b-wave. ERG recording was performed using a xenon white flash, 1000 Hz sampling frequency, 0.312 - 300 Hz cut off filter, 500 ms recording time, 10 ms baseline prior to flash, and no notch filter was used.

### Acoustic auditory brainstem responses (ABR)

Auditory sensitivity of the animals is evaluated by recording auditory brainstem responses to acoustic stimuli. Rabbits are anesthetized with xylazine and ketamine (and butorphanol to prolong anesthesia) and placed on a warm water-circulating heating in a sound attenuated chamber. Needle electrodes are placed under each pinna: test ear (reference) and contralateral ear (ground) and at the vertex (active) of the animal’s head to record the neural output. Tucker Davis Technologies (TDT) System III hardware and SigGen/BioSig software (TDT, Alachua, FL USA) is used to present the stimulus and record responses. Tones are delivered through an EC1 driver (TDT, aluminum-shielded enclosure made in house), with the speculum placed just inside the tragus. Stimulus presentation is 15 ms tone bursts, with 1 ms rise/fall times, presented 10 per second. Up to 1024 responses are averaged for each stimulus level. Responses are collected for stimulus levels in 10 dB steps at higher stimulus levels, with additional 5 dB steps near threshold. Thresholds are interpolated between the lowest stimulus level where a response was observed, and 5 dB lower, where no response is observed. ABR thresholds and suprathresholds will be tested at 3 or 4 frequencies (between 4 – 24 kHz). ABR sessions may last 45 - 60 min.

## Results

### Usherin is highly conserved in rabbit and human

The rabbit USH2A gene has two isoforms: (i) the short transcript containing 23 exons that encodes 1543 amino acids (Ensemble transcript ID: ENSOCUT00000014751.4); and (ii) the long transcript contains 74 exons encodes 5202 amino acids (NCBI Reference Sequence: XM_008268426.2). Analysis of protein sequences of usherin revealed that Usherin is highly conserved in human and rabbit, with 84% identify score for long isoforms and 85% identity score for short isoforms, and 91% positive score for both isoforms (**Fig.1A).** To our interest, as shown in **Fig. 1B**, rabbit exon12 of USH2A is of the same length of human exon13, both are 642 bp long, which notably is an in-frame length that is suitable for exon deletion-based therapy^39^. We examined the expression profiles of USH2A gene in rabbits by real-time PCR using two pairs of primers: (i) Pair 1 on exon11 and exon12, that detects both short and long isoforms; and (ii) Pair 2 on exon50 and exon51, that detects only the long isoform. As shown in Fig. **1C**, both isoforms of USH2A are exclusively expressed in the rabbit retina, but not in any other organs/tissues examined.

**Figure 1.**
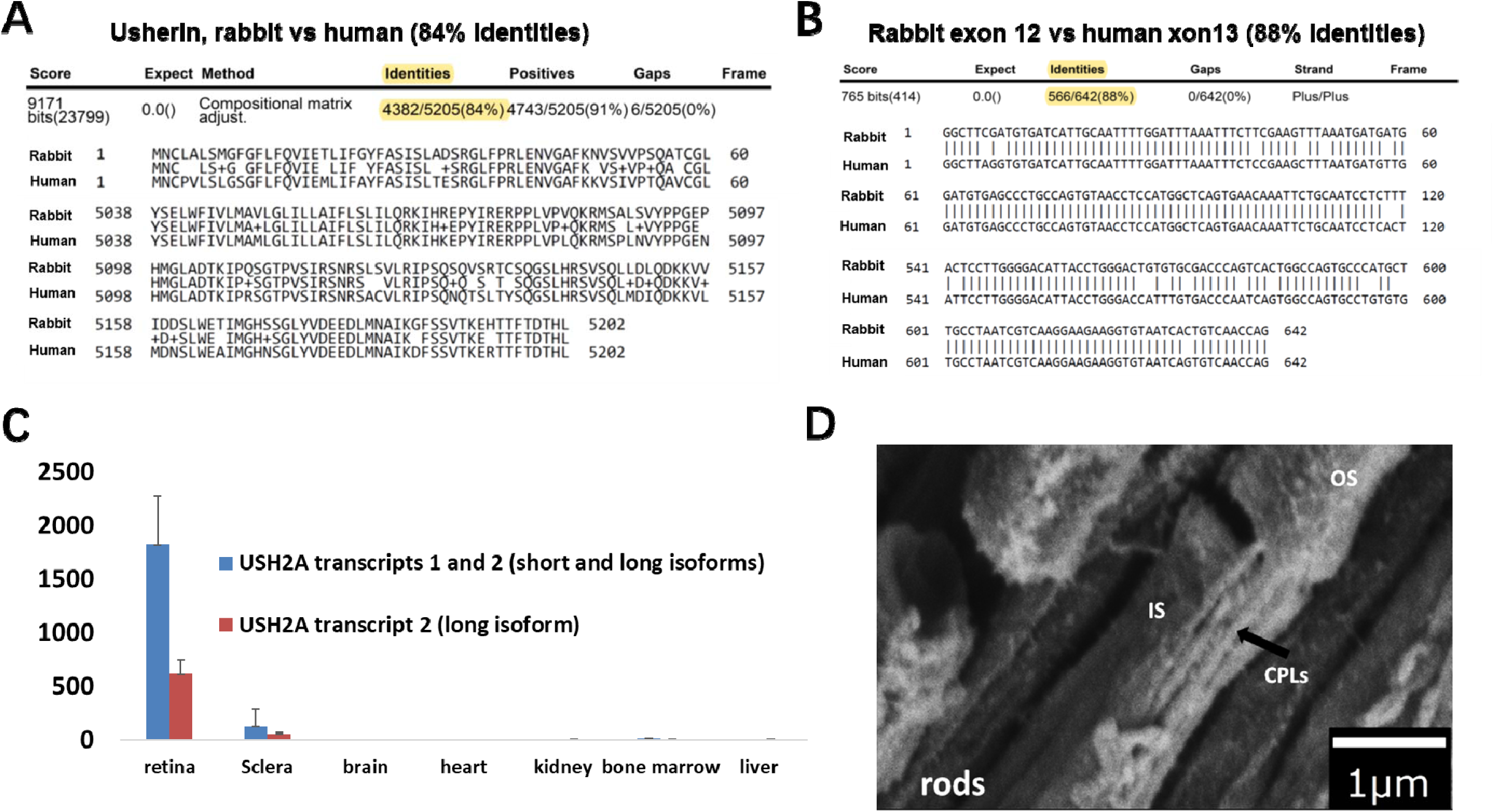
USH2A is conserved in rabbit and human. **(A).** Rabbit vs human Usherin protein long isoform sequence alignment using BlastP program. **(B).** Genomic DNA sequence alignment of rabbit exon 12 vs human exon 13 using BlastN program. **(C).** Real-time PCR detection of USH2A variants in adult rabbit organs. Primer pair 1(RTF1,RTR1) detects both variant 1 and variant 2 (v1+v2, blue bar); Primer pair 2(RTF2,RTR2) detects variant 2 only (orange bar). Values were normalized to18S rRNA expression. Y axes show the fold change relative to the expression level in brain. Error bars represent Standard Deviation. **(D).** Scanning electron microscopy shown Calyceal Processes like structures (CPLs) in rabbit photoreceptors. IS/OS: inner and outer segments of rod and cone photoreceptor cells.

### Calyceal processes like structures in rabbits

It has been reported that photoreceptor calyceal processes (CPs) are present in the retina of primates but absent from mice, suggesting that the presence/absence of CPs maybe the cause of the difference in visual phenotype between USH1 human patients and mice models^40^. It is possible that this structure (i.e. CPs) is also critical to the eye pathogenesis in USH2, as USH1 and USH2 proteins function together in higher order protein complexes^41^. Electron microscopy imaging results indicate that CP-like structures exist in rabbit retina **(Fig. 1D)**.

### Production of USH2A mutant rabbits

In efforts to model USH2 in rabbits, we chose to knock in the USH2A c.2299delG mutation, the most prevalent USH2A frameshift mutation found in human USH patients^5^, into rabbit genome. We designed four gRNAs targeting exon 12 of the rabbit USH2A gene (**Fig.2A**). All four sgRNAs could cut their target in test tubes efficiently with Cas9 protein (**Fig.2B**). T7E1 and Sanger sequencing assay showed that sgRNA1 achieved high efficiency of cleavage in rabbit embryos (**Fig.2B**). The guide RNA1 selected was co-introduced to rabbit embryos with Cas9 mRNA and a donor single stranded DNA harboring the intended c.2299delG mutation and 50 nucleotides homologue arms on each side. Totally 60 injected embryos were surgically transferred into the oviduct of two synchronized recipients. All 9 term kits were identified as USH2A mutant animals with 1 of them carrying 15.08% of the c.2299delG mutation detected by ear skin deep sequencing (**Fig.2 C&D, supplementary Fig.1**). these data demonstrate that USH2A mutant founder rabbits can be produced by CRISPR/Cas9 very efficiently. Sperm DNA analysis by targeted deep sequencing in founder that carries the c.2299delG mutation show a high percentage of presence of this mutant allele (10.22% HDR, **Fig.2D**). Among the knockout founders, one male (Founder#2) was mated with two wildtype rabbits producing 19 kits, 9 of which carried frameshift indels mutations predicted to cause premature stop codon (+14bp, -11bp, and -1 bp, **Fig.3B**). these data show that both knock-in and knockout USH2A rabbits are germline transmitting.

**Figure 2.**
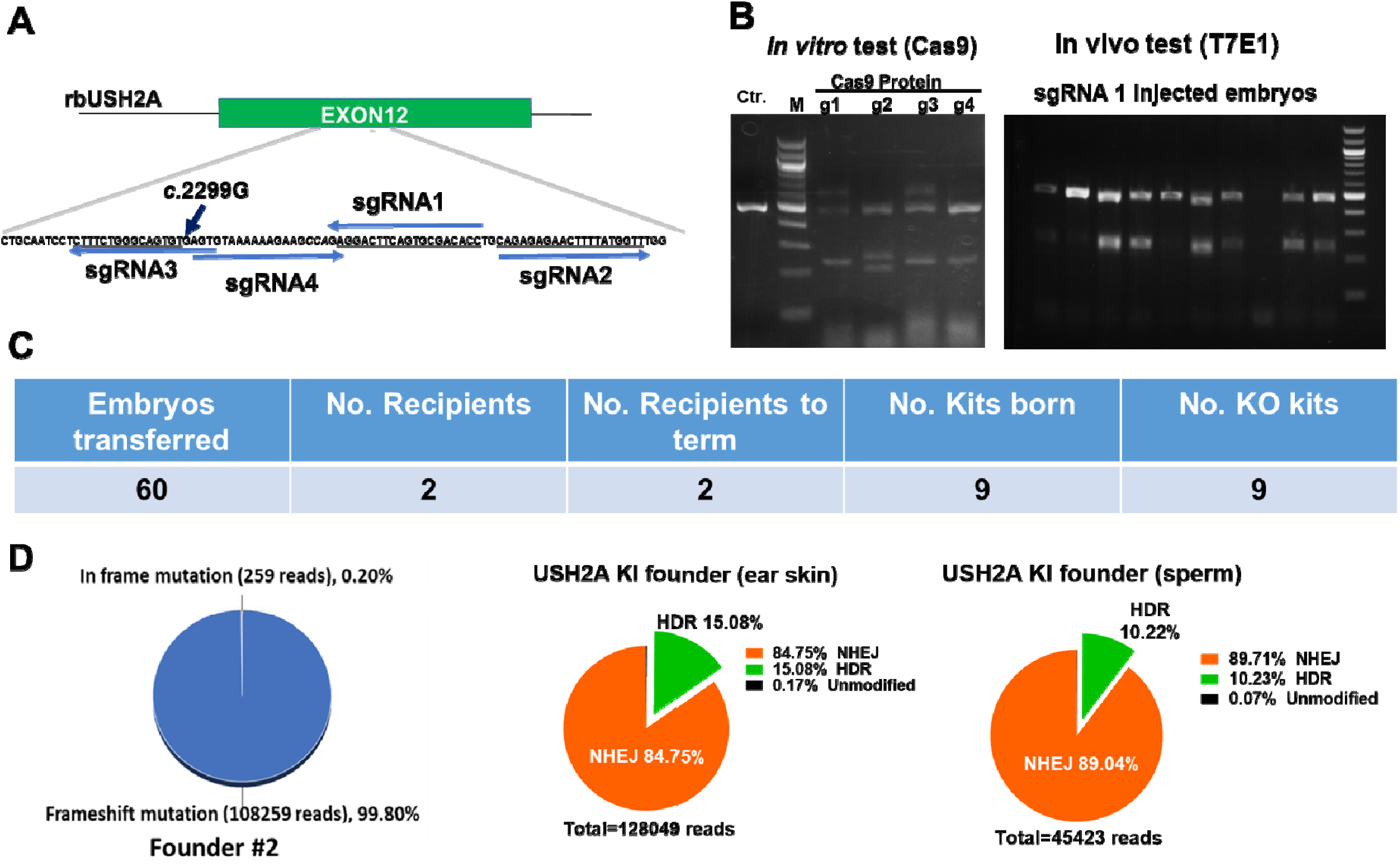
Production of USH2A mutant rabbits. **(A).** Illustration of CRISPR/Cas9 mediated targeting strategy to produce rbUSH2A mutant rabbits. **(B).** Four candidate sgRNA were designed and tested in vitro (left panel). sgRNA1 was validated in rabbit embryos using T7E1 analysis (right panel). In right panel, each lane representative one injected embryos. The PCR product of 479 bp will be cleaved into 238 bp and 241 bp bands if the embryos have indels generated at the gRNA target. M, NEB 100 bp DNA ladder. **(C).** Production of the USH2A mutant founder rabbits through embryo transfer. **(D).** Representative NGS analysis of the founder rabbits ear biopsy showed high frequencies of both indels (NHEJ) and knock-in (HDR) mutations (left and mid panel). The germline transmission of the mutations was confirmed by targeted deep sequencing in semen collected from the knock in founder.

**Figure 3.**
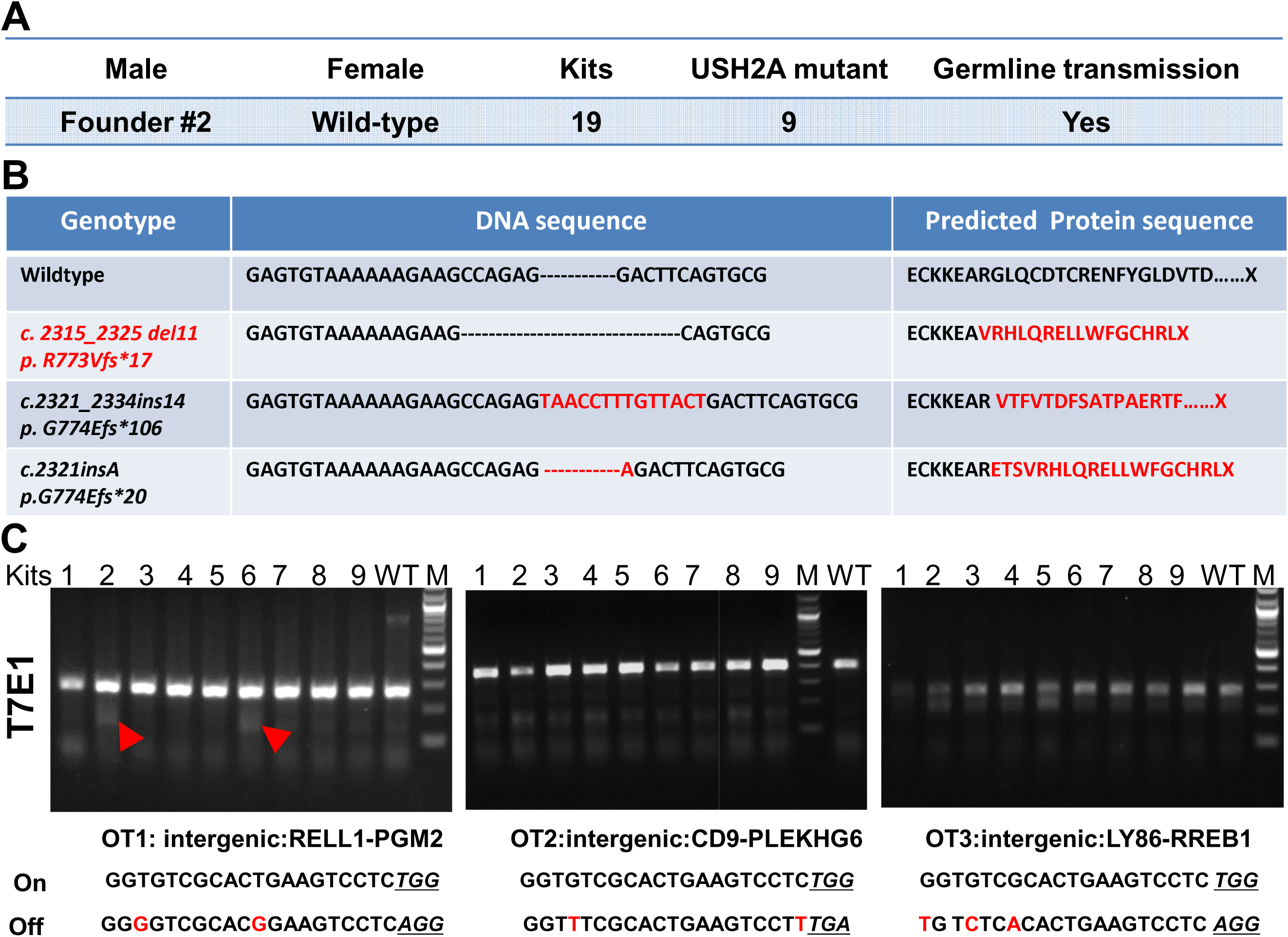
USH2A KO line establishment and off-target analysis. **(A).** Breeding of the founder male rabbit mated with two female wildtype rabbits. **(B).** Mutated USH2A DNA sequence and the predicted protein sequence found in the F1 generation USH2A KO rabbits. **(C).** Detection of off-target indels using T7E1 analysis in F1 generation USH2A mutant rabbits with a wild-type rabbit as control. Red arrow heads showing the indels detected at off-target 1 locus in F1 generation kits #2 and #6. M, NEB 100 bp ladder DNA marker; On, on-target sequences; Off, off-target sequences; Nucleotides in red color indicate the mismatches of the gRNA and the potential off-target sequence. NGG/NGA PAMs were highlighted by underlines.

### Off target analysis

To test the off-target effects, we chose the top three potential off-target sites predicted by the gRNA design software to test off-target effects, in which the predicted off-target sites were PCR amplified and analyzed with T7E1 assay and confirmed by Sanger sequencing. No indels were detected at OT2 and OT3 in any of the nine F1 rabbits (**Fig. 3C**), whereas at off-target1 (OT1), two of the nine F1 generation rabbits showed indels (GCTG to CTC) that are located 32 bp away from the predicted Cas9 cleavage site (**Fig.3C, Supplementary Fig.2**). As this region is an intergenic region, it is more likely a natural polymorphism. Nevertheless, to avoid the potential adverse effects, these two F1 rabbits carrying the OT1 site indels were excluded from the breeding program.

### Exon 12 mutation in rabbits result in nonsense-mediated mRNA decay

Upon sexual maturation, one of the F1 rabbit carrying del11 mutation was used to establish the USH2A KO (USH2A-/-) line. This mutation is predicted to cause a premature stop codon, which may lead to the nonsense-mediated mRNA decay and the production of non-functional truncated Usherin. Real time PCR detection of the USH2A transcripts in the retina tissue isolated from a founder animal carrying high percentage of indels mutation (Founder #2 in **Fig.2D.)** and a USH2A KO homozygous rabbit. As shown in **Figure 4**, the expression of USH2A transcripts was decreased more than 50% in the founder animal retina compared with wildtype controls, and the expression level further decreased to less than 30% that of wildtype controls in the USH2A KO retina, indicate nonsense-mediated mRNA decay did happened in the USH2A KO rabbits. We expect that the introduced mutation results in decreased mRNA level and the residual mRNA carrying the mutation leading to premature termination of usherin translation.

**Figure 4.**
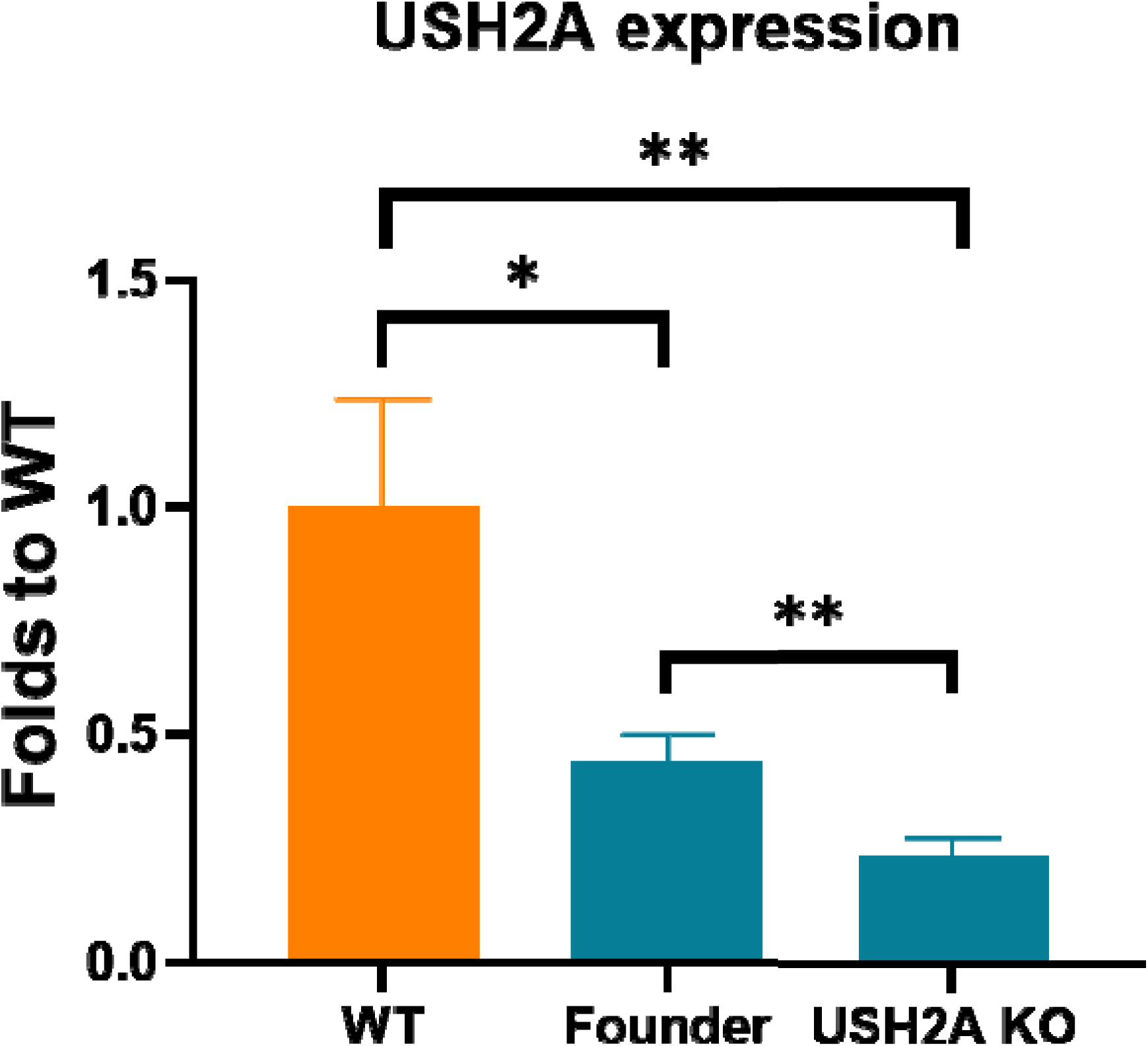
USH2A expression in USH2A KO rabbit retina. Real-time PCR detection of USH2A variants in retina tissue of a USH2A KO rabbit using the primer pair 1 in Fig.1, which detects both long and short variants of USH2A. Values were normalized to GAPDH RNA expression. Y axes show the fold change relative to the expression level in wild type controls (3 animals). Error bars represent Standard Deviation.

### Eye phenotype of the USH2A rabbits

As shown in **Figure 5A**, USH2A KO rabbits had normal retinal and choroidal vasculature. In addition, ophthalmic examination confirmed that all USH2A rabbits had ophthalmoscopically normal and healthy corneas, anterior chambers, and clear lenses. There was no difference in the fundus appearance between WT and USH2A KO rabbits. It should be noted that these USH2A KO animals were on an albino background, and the characteristic bone spicule pigmentation of the retina seen in RP eyes would therefore not be expected. FA and ICGA imaging shows normal retinal and choroidal vascular morphology. In contrast, hyper-FAF spots were detected in the retina of the USH2A KO rabbits as early as 4 months old. OCT images also indicated changes at the photoreceptor layer with hyper-reflective foci at the level of the photoreceptor IS and OS segments as early as 4 months old.

**Figure 5.**
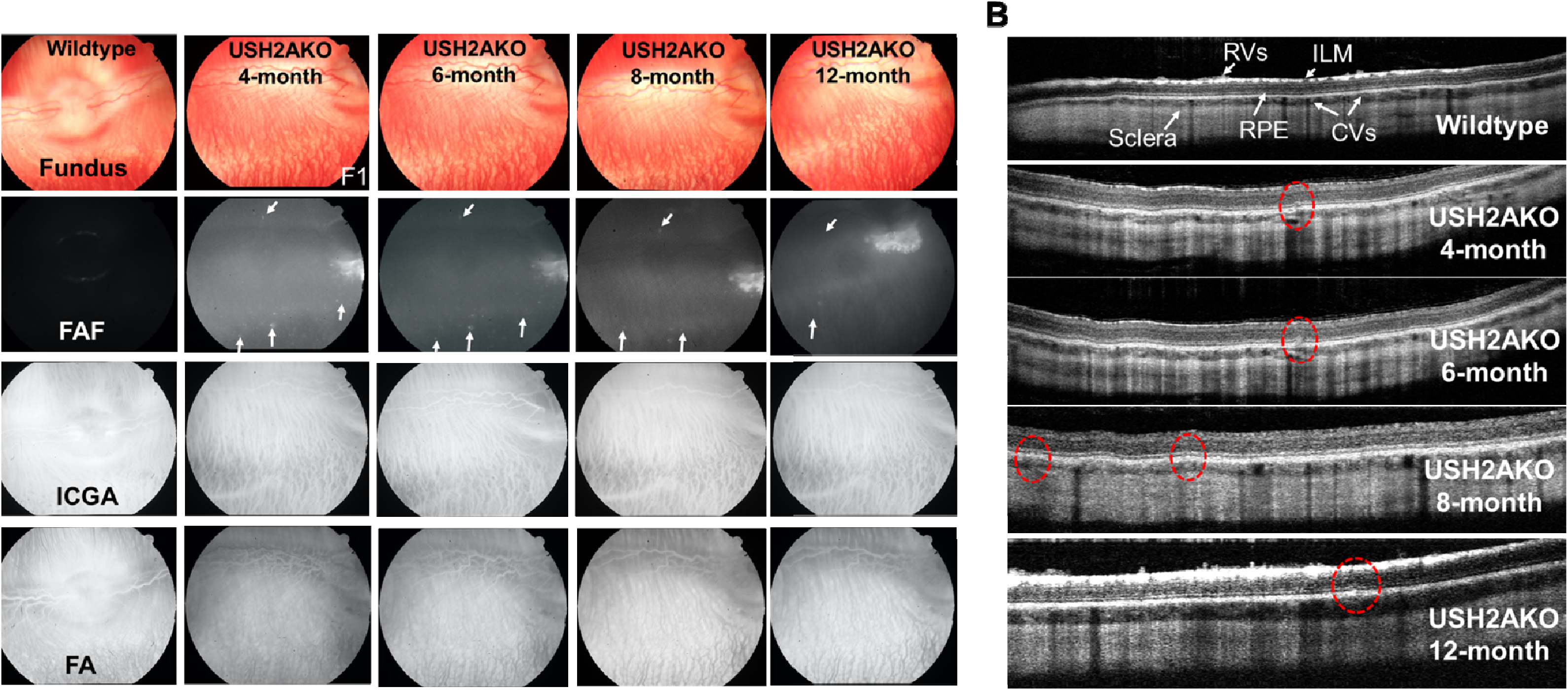
Retinal imaging in USH2A KO rabbit. **A).** Fundus photographs (Fundus), fundus autofluorescence (FAF), indocyanine green angiography (ICGA), and fluorescein angiography (FA) images obtained from a USH2A KO rabbit at 4-month-old to 12-month-old demonstrating hyper-FAF spots (arrows). **B)**. Spectral domain optical coherence tomography (OCT) images of a USH2A KO rabbit at 4-month-old to 12-month-old demonstrating hyper-reflective foci at the level of the photoreceptor IS and OS segments in the photoreceptor layer (red dotted circles).

OCT imaging was performed on both WT and USH2A KO rabbits as illustrated in Figure 5B. B-scan OCT image obtained from WT rabbits show normal and healthy retina with the different retinal layers such as the nuclear layers, plexiform layers, photoreceptors, retinal vessels (RVs), choroidal vessels (CVs), RPE, inner limiting membrane, and sclera. No evidence of photoreceptor or RPE disruption or damaged was observed in WT rabbits. On the other hand, hyper-reflective foci were noted at the level of the photoreceptor inner and outer segments and RPE in USH2A rabbits at 4 months old as marked by red dotted circle, which increased over time with more disruptions at 8 and 12 months old.

### Loss of retinal function in USH2A KO rabbits

To investigate whether the disruption of USH2A gene causes retinal abnormalities, visual function of USH2A KO and WT control rabbits were compared by ERG analyses at different ages under scotopic and photopic conditions. At age 7 months, the USH2A KO rabbits displayed rod response (scotopic 24 dB), scotopic combined rod and cone response (scotopic 0dB), and photopic response b-wave amplitudes (**Fig.6A,B Fig. S3**) and 32 Hz flicker amplitudes (**Fig. 6C,D**) significantly lower than WT counterparts. The responses of USH2A KO rabbits were 10% to 20% lower than those of WT control (P=0.0456 and 0.042 respectively by t-test) as shown in **Figures. 6B, 6D** and **Fig. S3**. At age 15-22 months, scotopic b-wave amplitudes and 32 Hz flicker amplitudes became further reduced with the responses of USH2A KO more than 50% lower than those of WT controls (P =0.0073 and 0.0031 respectively by t-test). This age dependent decline suggests progressively loss of photoreceptor function in USH2A KO rabbits. Implicit time did not show a significant difference between WT and USH2A KO over time (**Fig. S4**).

**Figure 6.**
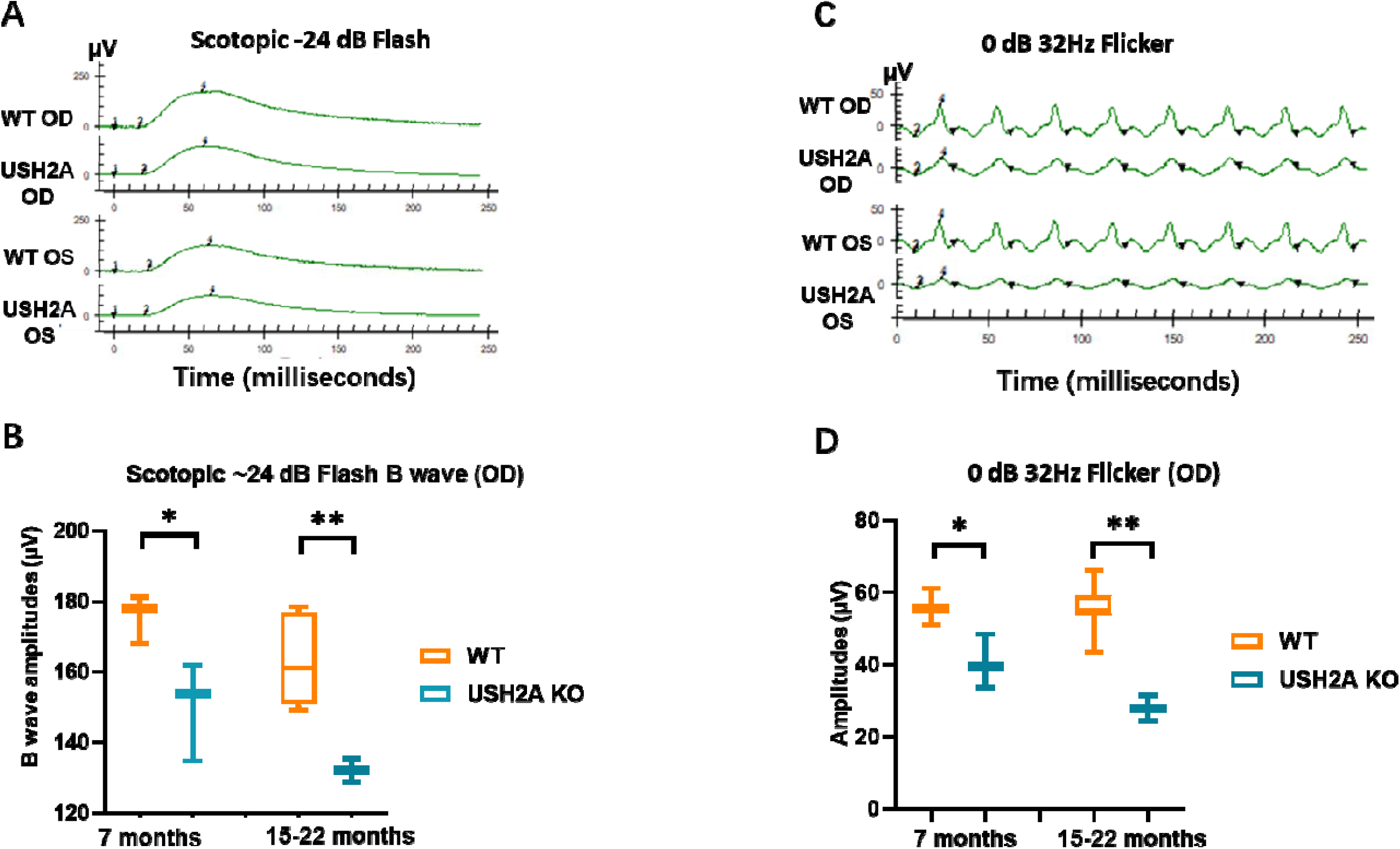
Progressive retinal degeneration in USH2A KO rabbits. **A).** Representative rod ERGs recorded at scotopic -24 dB flash from a 15-month-old USH2A KO rabbit with an age matched wild-type rabbit as Control. **B).** Full field electroretinography (ERG) demonstrated a significant reduction in rod response amplitude at 7 months that is further reduced at 15-22 months in USH2A KO rabbits compared with age matched wildtype (WT) rabbits. **C).** Representative ERGs recorded at 0 dB 32Hz flicker demonstrating cone response from a 15-month-old USH2A KO rabbit with an age matched wild-type rabbit as control. **D).** A significant reduction in amplitude of cone ERGs by 7 months that is consistent and further reduced at 15-22 months were recorded in USH2A KO rabbits compared with age matched wildtype (WT) rabbits. ICGA: indocyanine green angiography, FA: fluorescein angiography, RVs: Retinal vessels. ILM: Inner Limiting Membrane. RPE: Retinal pigment epithelium. CVs: Choroidal Vessels. OD: Right eye; OS: Left eye. Data analyzed by unpaired t-test, * p<0.05, ** p<0.01.

### USH2A rabbit histopathology showed reduced photoreceptor nuclei in the ONL

To look for photoreceptor degeneration characteristics of Usher-associated retinitis pigmentosa, the number of photoreceptor nuclei in the outer nuclear layer (ONL), which represents the rod and cone cell bodies, of a 16 months USH2A KO rabbit was compared with age matched control rabbits. Figure **6A** shows the H&E image of WT (left) and USH2A KO (right). These H&E images clearly show thinner ONL layer in the USH2A KO rabbit retina compared with the age matched wildtype control. **Figure 7B** represent the overview of a rabbit retinal section illustrating the locations of optic nerve (Myelinated region) and visual streak as well as the 10 spots for ONL nuclear number counting in **Figure 7C.** We found that the USH2A KO rabbit has reduced photoreceptor nuclear numbers throughout the whole retina compared to the age matched controls, indicating photoreceptor cell degeneration which explained the reduced ERG signals in **Figure 6.**

**Figure 7.**
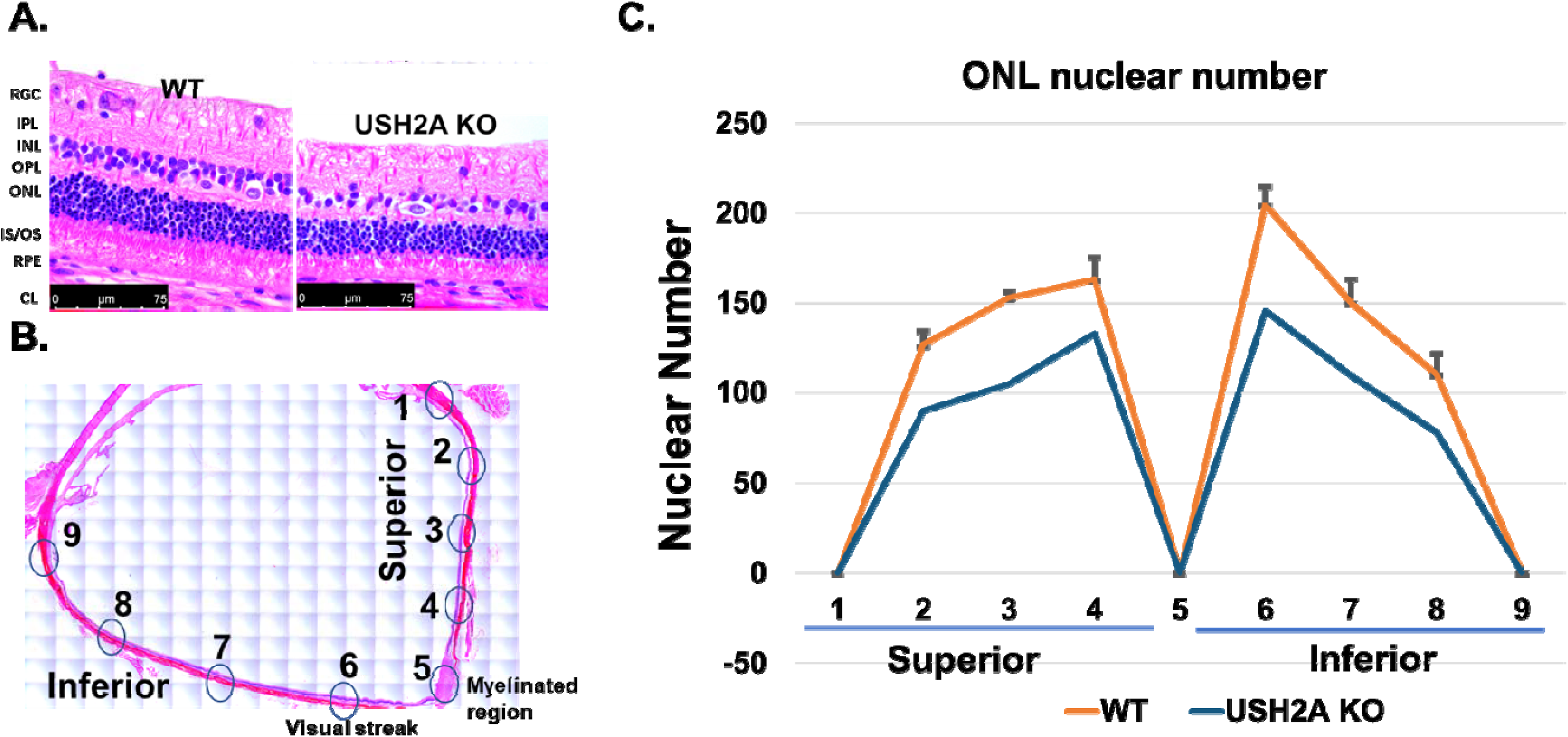
Histopathology analysis of USH2A KO rabbit retina. **A).** Representative retinal section prepared from a 16 months USH2A KO rabbit stained with H&E show reduced retinal layer thickness compared with age-matched wildtype (WT). Scale bars 75 µm. **B).** overview of a rabbit retinal section illustrating the locations for ONL nuclear number counting in Panel C. **C).** Counting of ONL layer nuclei numbers in a defined field of view (150 µm along the retina layer) at different locations on the retina indicated in panel B. Error bars represent SD. RGC: Retinal Ganglion cell layer; IPL: Inner plexiform layer; INL: Inner nuclear layer; OPL: Outer plexiform layer; ONL: Outer nuclear layer; IS/OS: Inner and Outer segments of rod and cone photoreceptor cells; RPE: Retinal pigment epithelium; CL: Choroid layer.

**Figure 8.**
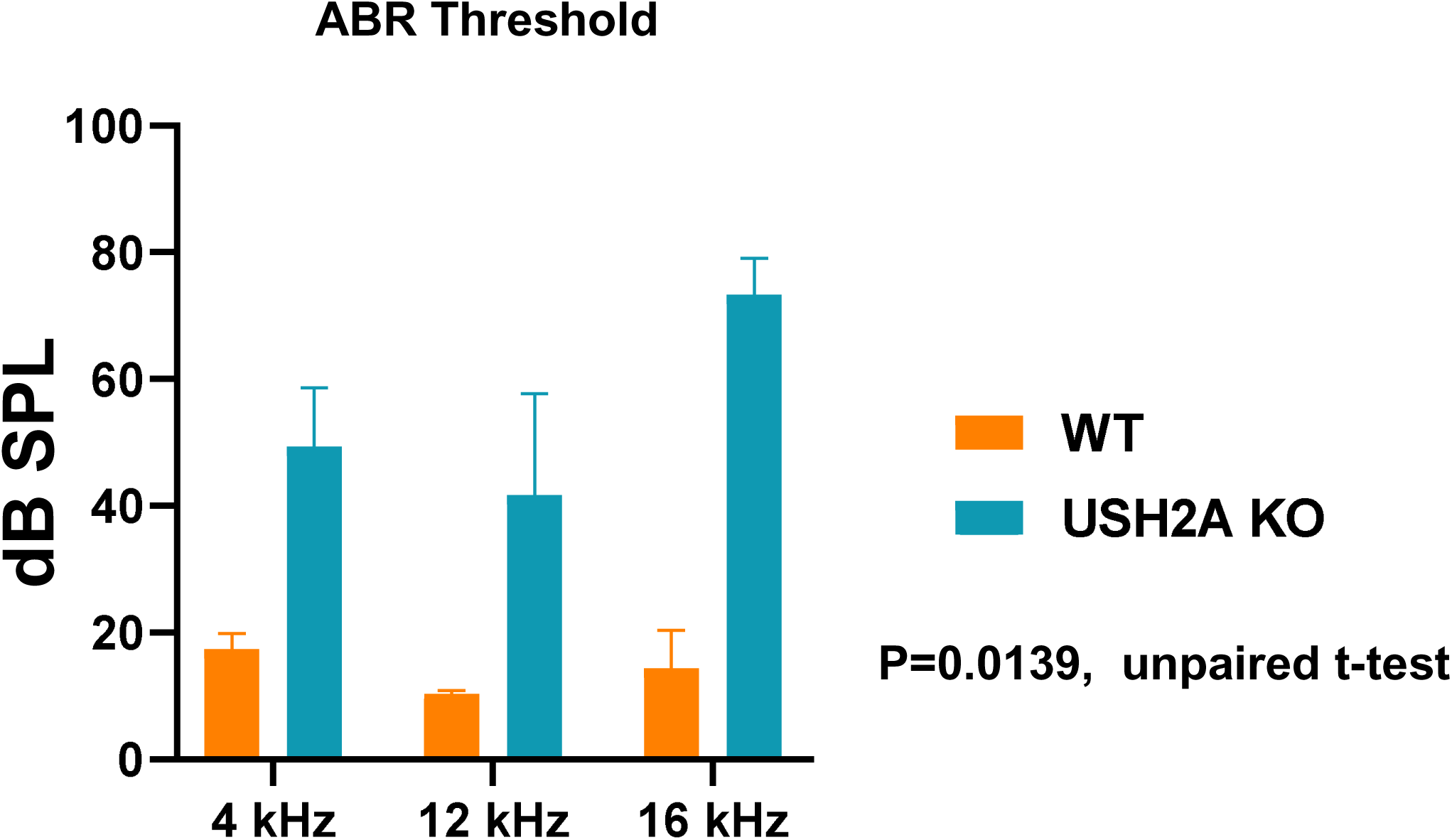
Hearing loss in USH2A KO rabbits. Auditory brainstem response (ABR) test demonstrating significant hearing loss in the USH2A KO rabbits at 5 months old compared with wildtype (WT) control rabbits (p=0.0139, n=3).

### USH2A KO rabbits showed moderate to severe hearing loss

USH2 affects both hearing and vision. In this study, auditory brainstem response (ABR) was tested in three USH2A KO rabbits at 5 months old. As shown in figure 7, ABR threshold at three different frequencies tested (4 kHz, 12 kHz, and 16 kHz) were all increased by 2-3 times in the USH2A KO rabbits compared with wildtype control rabbits (p=0.0465, n=3) **(Fig. 7)**, indicating moderate to severe hearing loss in these rabbits.

## Discussion

Usher syndrome type 2 is characterized by hearing loss and early adulthood-onset of RP. Retinal degeneration in patients is apparent by fundus examination and the progressive reduction in ERG amplitudes over the course of the disorder. A targeted Ush2a knock-out mouse demonstrates only mild retinal degeneration with late age of onset^6^. A spontaneous mutant mouse model, Kunming, shows a rapid, early-onset retinal degeneration, but contains mutations in two genes known to be involved in inherited retinal dystrophies: Ush2a and Pde6b ^47^. Recently, bright light induction has been reported to be able to induce the damage of the rod photoreceptors in several of the USH1 and USH2 mouse models. However, these experiments were in a 129 Sv/j background, which is inherently more sensitive to light-induced photoreceptor cell damage ^48, 49^. Overall, these models only showed slightly reduced rod function indicated by reduced scotopic b wave amplitude, but no cone function reduction was detected. This is probably due to their late onset features, as cones are affected later in the disease course compared with rods. Due to the short life span of the mice, it is very difficult to study USH2 disease in mouse models. Here, we have successfully established USH2A KO rabbit line. We found that targeted disruption of the USH2A gene in rabbits resulted in hearing loss and severe retinal degeneration starting as early as 4 months of age, which is equivalent to human adolescence, and showed progressive retinal degeneration mimicking the RP in USH2 patients. To our knowledge, it is the first genetic preclinical large animal model that manifest eye phenotype of USH2.

In humans, the location of highest acuity in the retina is a circular area termed the fovea which boasts the highest concentration of cones, the lowest concentration of rods, and much smaller receptive field sizes for all cells. The area of greatest acuity in rabbit retina is not a single point, but rather an elongated “streak” running across the retina. The present of the visual streak made rabbits very useful for studying retinal degenerative diseases in contrast to mice. As shown in figure 6, the reduction of the ONL nuclear numbers in USH2A KO rabbits were more apparent in the visual streak area (point 6), suggesting loss of cone photoreceptors. It is reported that severe visual phenotype seen in syndromic USH2A patients compared with the non-syndromic USH2A patients could relate to a greater extent of cone dysfunction indicated by significantly reduced 30Hz-ficker ERG amplitudes^50^. In this study, in addition to the reduced rod function detected by the reduced scotopic b wave amplitudes in ERG, our USH2A KO rabbit models also showed significantly reduced 32Hz-ficker ERG amplitudes, which mimicking the cone function reduction in human patients.

In conclusion, we have succeeded in generating a rabbit model of USH2. Although further studies are needed to fully characterize the natural history of hearing loss and retinal degeneration phenotype in USH2A KO rabbits and to determine the exact mechanism of photoreceptor dysfunction observed in this model, we believe that this USH2A mutant rabbit model will serve as a useful large animal model with which to study the pathophysiology of RP in USH and develop novel treatments. The successful replication of the RP phenotype in USH2A rabbit models in this study as well as the extension of gene targeting technology to rabbits by CRISPR/Cas9 technology motivating efforts to develop rabbit models for other types of Usher Syndrome as well as other hereditary retinal diseases.

## Acknowledgements

This work was supported by National Institutes of Health grants, R01-HL147527 (YEC), R21 GM140359-01 (DY), K08EY027458 (YMP), Alcon Research Institute Young Investigator Grant (YMP), unrestricted departmental support from Research to Prevent Blindness, and the University of Michigan Department of Ophthalmology and Visual Sciences. This research utilized the Core Center for Vision Research funded by the National Eye Institute (P30 EY007003).

## Author contributions statement

D.Y, Y.M.P., Y.P and Y.E.C conceived the experiments. D.Y, Y.M.P, V.P. N, J.S, Y. L, D.P, D.D, J.X, and J.Z, conducted the experiments. D.Y, Y.M.P, K.T.J and Y.E.C analyzed the results and wrote the manuscript. All authors critically reviewed the manuscript.

## Additional information

Competing financial interests: The authors declare no competing financial interests.

---

**Supplementary Figure 1.**
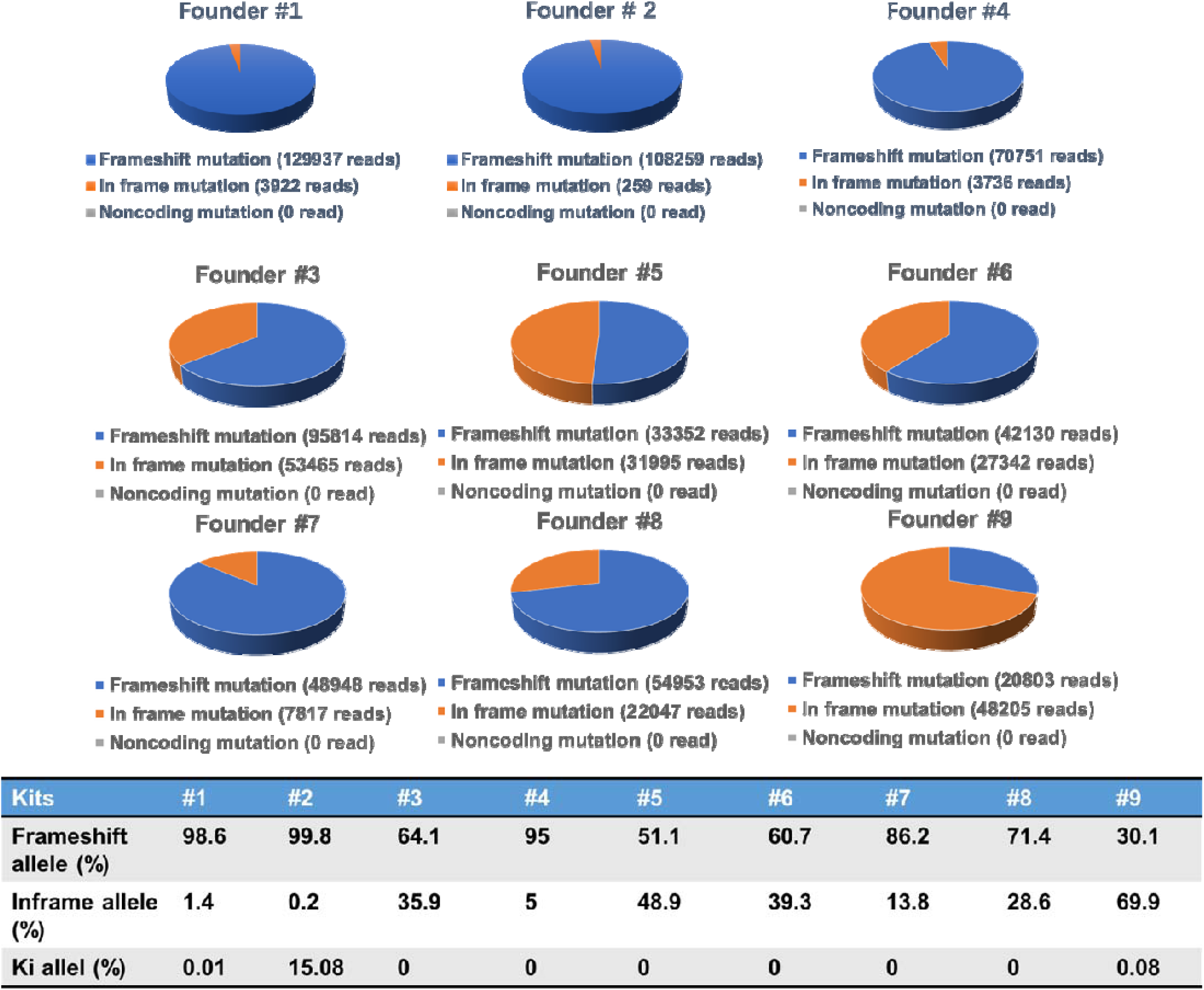
NGS sequencing of USH2A KO founder rabbits.

**Supplementary Figure 2.**
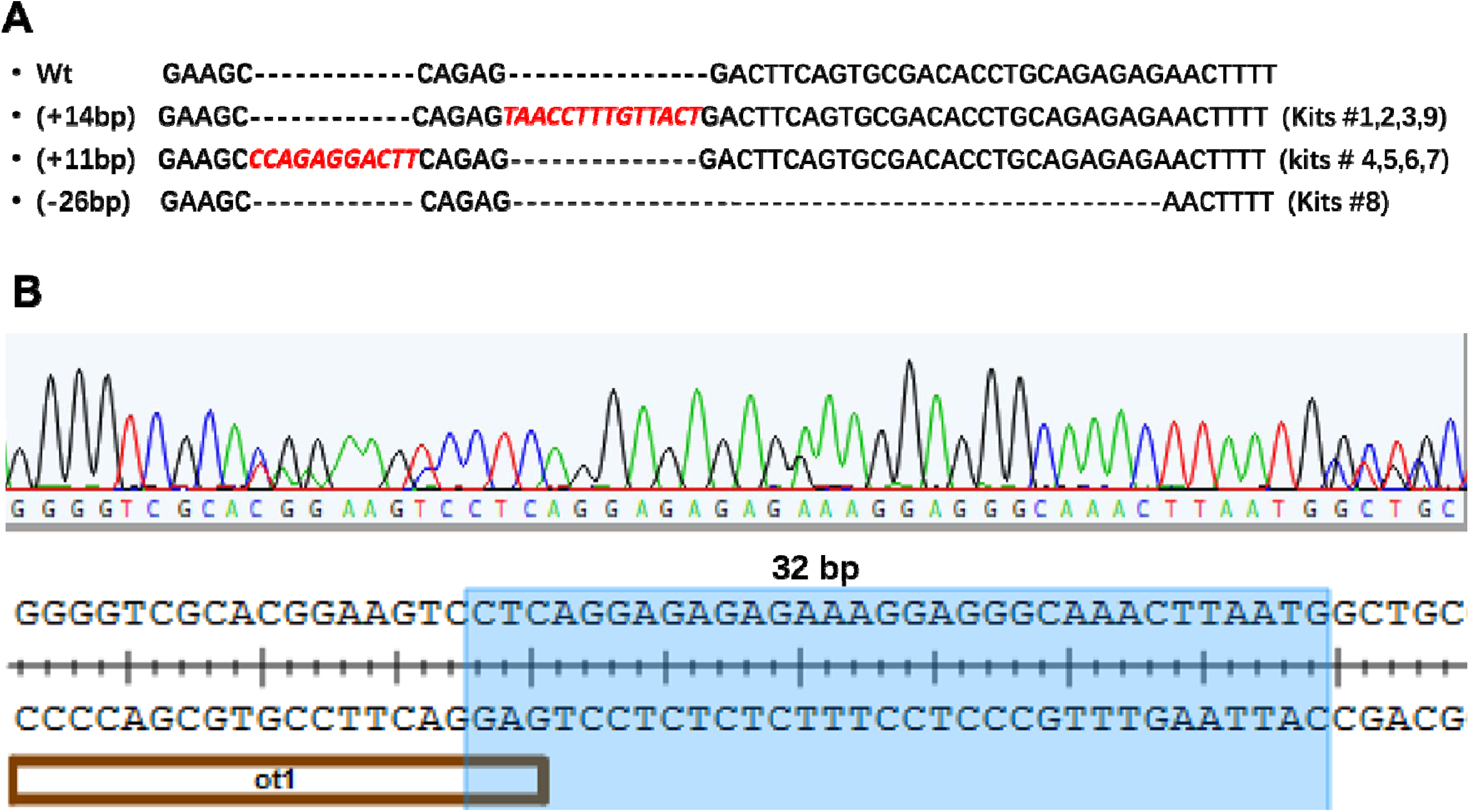
Germline transmission of the mutant alleles in F1 generation rabbits. **(A)** and Sanger sequencing of the OT1 off-target mutation **(B)**. Ot1: the off-target binding sequence. The 32 bp shadow part indicated the distance from the indels mutation (double peaks) site to the Cas9 cleavage site (3 bp upstream of the AGG PAM sequence)

**Supplementary Figure 3.**
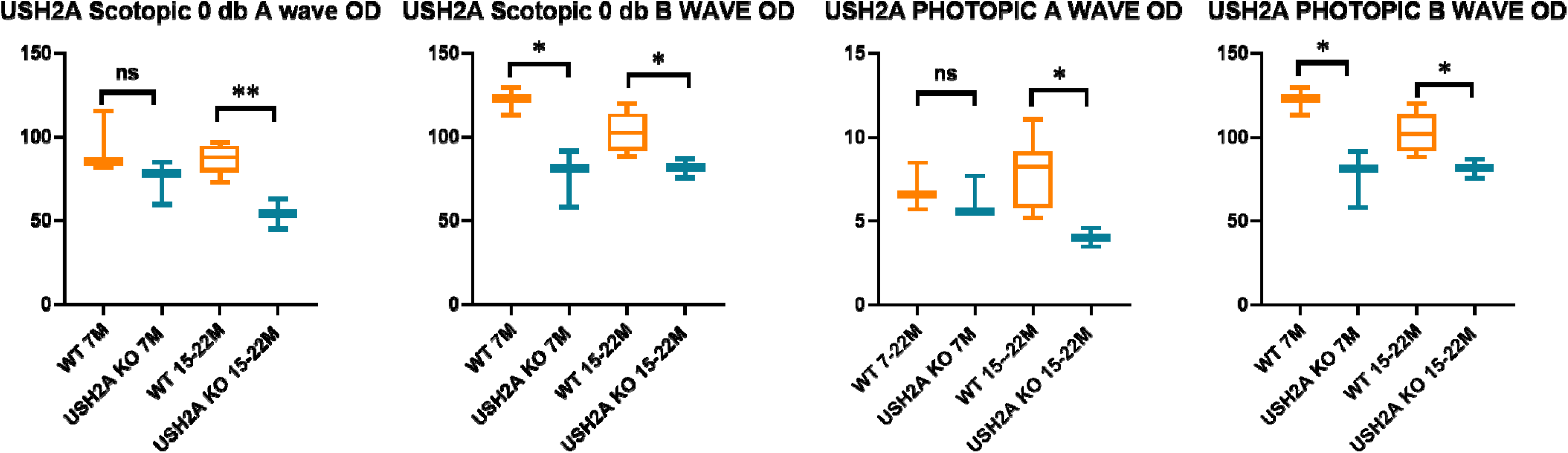
ERG Amplitudes in USH2A KO rabbits.

**Supplementary Figure 4.**
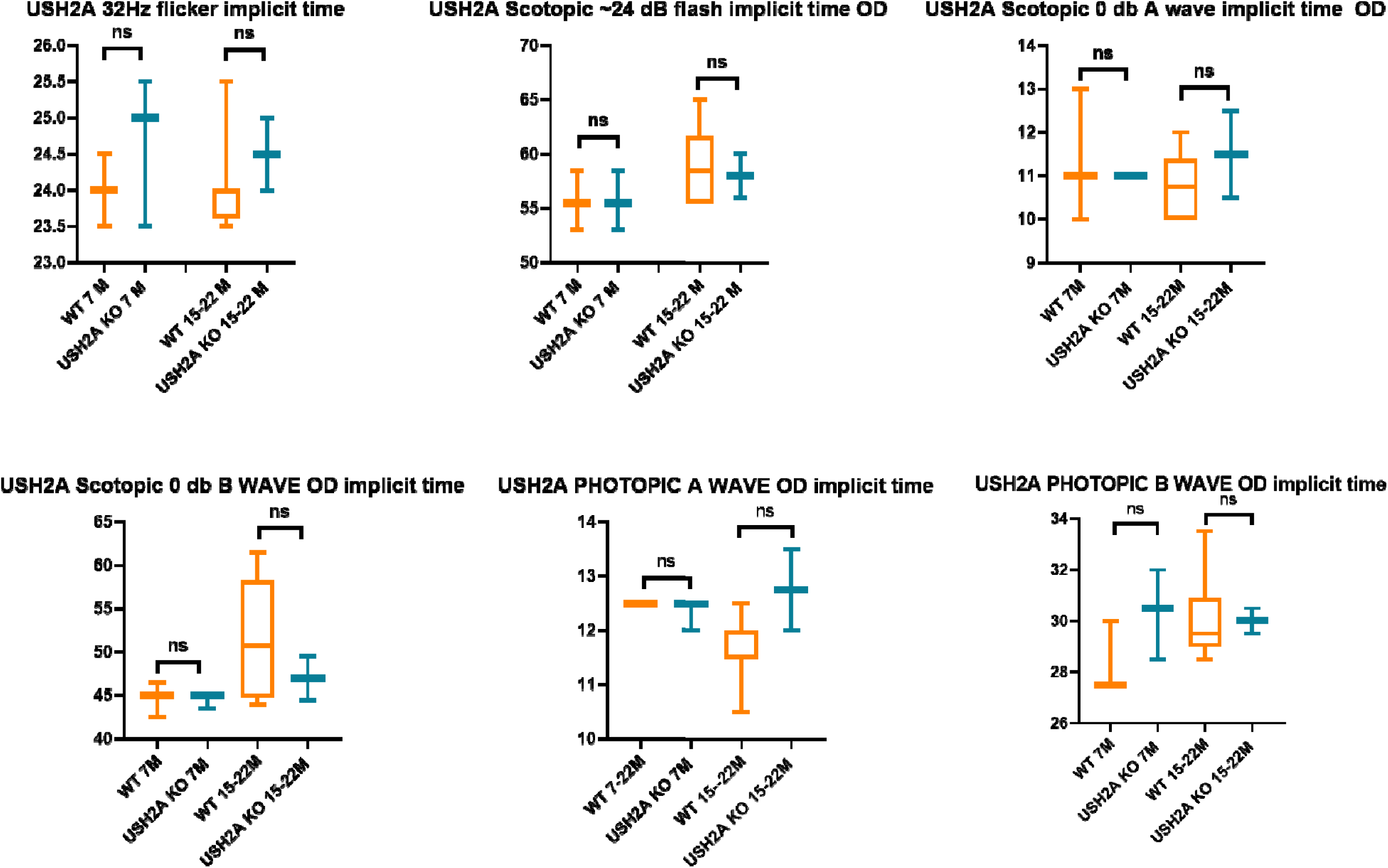
ERG implicit time in USH2A KO rabbits.

**Supplementary table 1.**
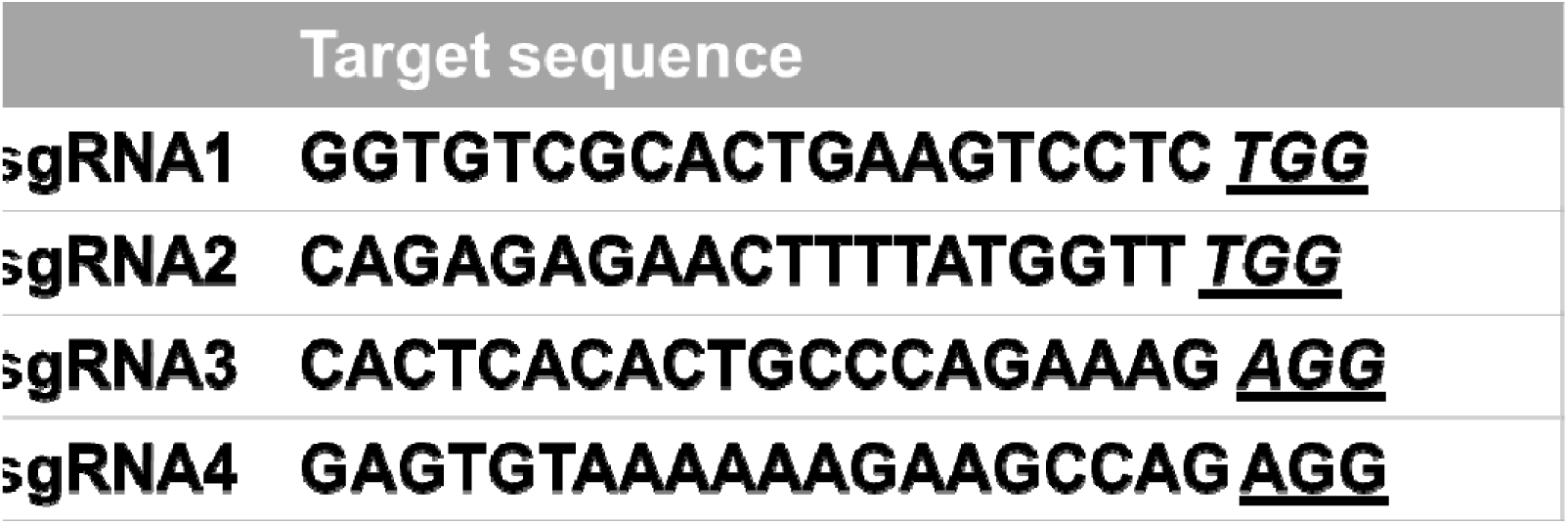

**Supplementary table 2.**
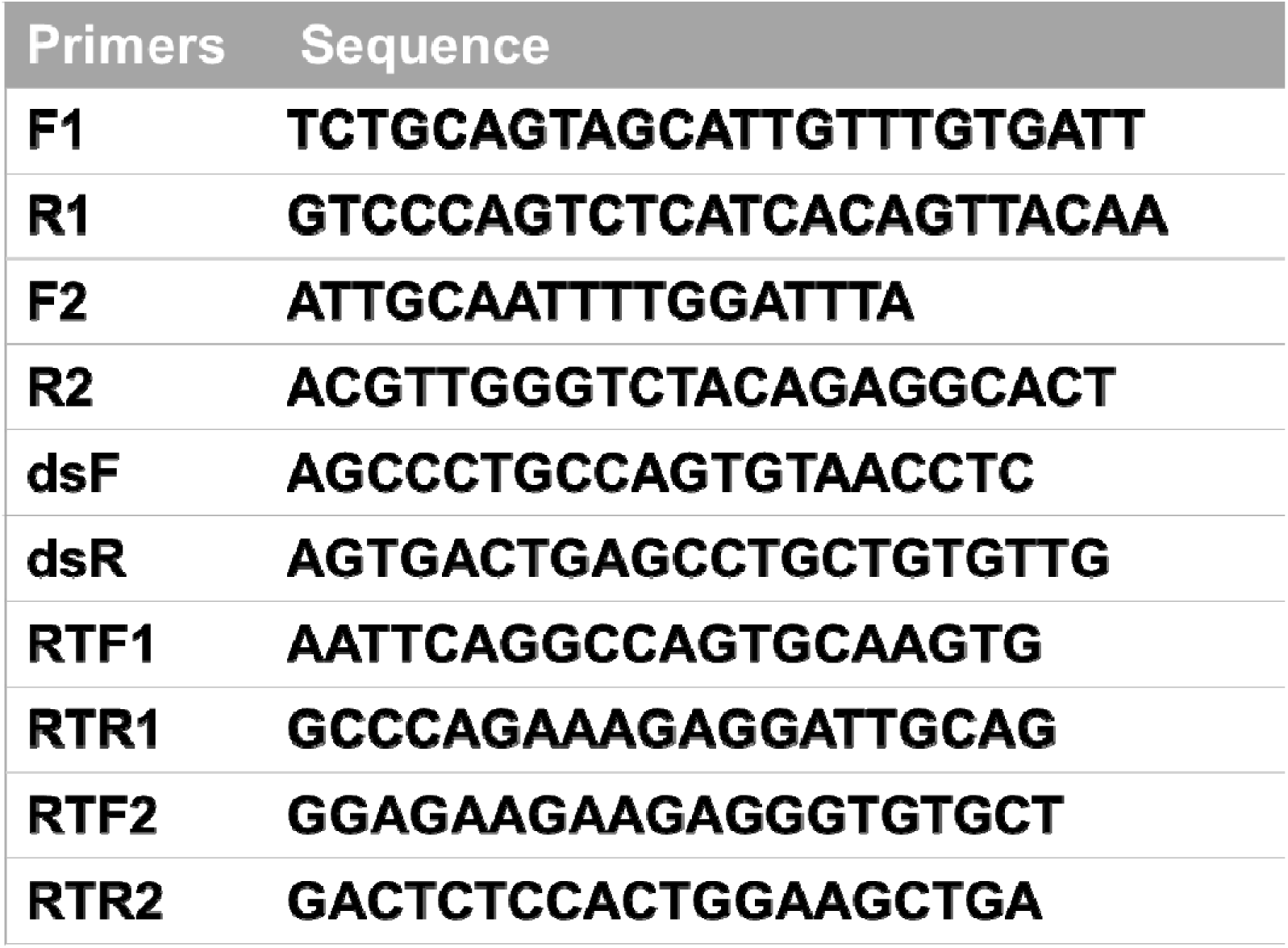

## Notes

### Competing Interest Statement

The authors have declared no competing interest.

